# Comparison Between Normobaric Hypoxia Altitude Simulation Test and Altitude Hypoxia Predictive Equations in Cystic Fibrosis Patients

**DOI:** 10.1101/570754

**Authors:** Christine Costa, André Barros, João Valença Rodrigues, Richard Staats, Mariana Alves, Pilar Cardim, Carlos Lopes, Cristina Barbara, Luís F. Moita, Susana Moreira

**Author notes:** Corresponding author: Christine Costa, Luís Moita or Susana Moreira.

## Abstract

**Background:** The Hypoxia Altitude Simulation Test (HAST) is the *Gold Standard* to evaluate hypoxia in response to altitude and to decide on in-flight requirements for oxygen supplementation. Several equations are available to predict PaO_2_ in altitude (PaO_2alt_), but it remains unclear whether their predictive value is equivalent. We aimed to compare the results obtained by the available methods in a population of cystic fibrosis (CF) adults.

**Methods:** Eighty-eight adults (58 healthy controls and 30 CF patients) performed a spirometry followed by an HAST. HAST results were compared with the predicted PaO_2alt_ made by five equations: 1^st^: PaO_2alt_= 0,410 x PaO_2ground_ + 1,7652; 2^nd^: PaO_2alt_= 0,519 x PaO_2ground_ + 11,855 x FEV_1_ (L) − 1,760; 3^rd^: PaO_2alt_= 0,453 x PaO_2ground_ + 0,386 x FEV_1_ (%) + 2,44; 4^th^: PaO_2alt_= 0,88 + 0,68 x PaO_2ground_; 5^th^: PaO_2alt_= PaO_2ground_ − 26,6.

**Results:** None of the controls required in-flight oxygen neither by HAST or by the five predictive equations. Eleven CF-patients had PaO_2alt_ < 50 mmHg, accessed by HAST. The positive predictive value was 50% (1^st^), 87.5% (2^nd^ and 3^rd^), 77.78% (4^th^) and 58.33% (5^th^). Areas under the curve were 78.95% (1^st^), 84.69% (2^nd^), 88.04% (3^rd^) and 78.95% (4^th^ and 5^th^). FEV_1_ and PaO_2ground_ were correlated with HAST results.

**Conclusions:** The 3rd equation gave the best predictions in comparison with results obtained by HAST. However, because the individual differences found were substantial for all equations, we still recommend performing a HAST whenever possible to confidently access in-flight hypoxia and the need for oxygen.

## Introduction

The quality of life and life expectancy of patients with cystic fibrosis (CF) have increased substantially during the past years^1^ due to medical improvement and preventive initiatives, which currently enable these patients to participate in activities that were not previously feasible, such as tourism.^2^ As a result, physicians are increasingly requested to evaluate these patients for their ability to fly safely and the need for oxygen supplementation.^3,4^ Commercial flights expose individuals to a lower atmospheric pressure and consequently to a lower oxygen content,^2^ as passenger cabins of commercial aircraft at maximal cruising altitude are pressurized to an altitude equivalent of 2438 m (8000 feet),^1,3,5-7^ which is equivalent to breathing 15% oxygen at sea level.^3-10^ This altitude hypobaric hypoxia can be tolerated in healthy individuals, but may cause severe hypoxemia in patients with pulmonary disease.^3,10-14^ CF-patients are therefore prone to a substantial and unsafe reduction in the partial arterial oxygen pressure (PaO_2_) during the flight^1,12^ that may lead to a severe respiratory decompensation.^15^

The Aerospace Medical Association^2^ and British Thoracic Society (BTS)^4^ recommend to perform a hypoxic altitude simulation test (HAST) to assess whether patients need in-flight oxygen supplementation. The HAST is considered the gold standard^11^ and it can be done by artificially reducing inspired oxygen to similar levels as those experienced at 2438 m (8000 feet) for 20 min by either reducing the fraction of inspired oxygen to 15%^1,4,7,11^ or by reducing atmospheric pressure to 565 Torr (75 kPa) in a hypobaric chamber.^4,11^ The normobaric HAST is usually the preferred technique, as it is more accessible and inexpensive than the HAST performed in a hypobaric chamber. The expected PaO_2_ (PaO_2HAST_) is then determined from measurements of arterial blood gases (ABG).^11^ A subject is judged to require in-flight oxygen supplementation if the PaO_2HAST_ falls below 50 mmHg, although this arbitrary cut-off value has little supporting evidence.^4^

Security cut-offs have been proposed that can be used when HAST is not available. In the case of CF, patients with FEV_1_ > 50% or PaO_2ground_ > 60 mmHg can safely be allowed to travel without oxygen supplementation.^3,12^ In addition, several investigators have developed predictive equations that estimate PaO_2_ at altitude (PaO_2alt_) using measurements made at sea level.^6,11,14,16,17^ The BTS guidelines refers to four of them, but they have been derived almost exclusively from patients with chronic obstructive pulmonary disease who undergone a HAST.^4^ Only one study, performed by Kamin et al.^17^ (in our study referred as 5^th^ equation) testing 12 adults with CF with mild to moderate respiratory insufficiency (mean PaO_2_: 79 mmHg) proposed one equation to predict the expected hypoxemia for flights of up to 3,5 hours in duration.^17^

The aim of this study was to compare HAST results with those predicted by the available equations in a population of adults diagnosed with CF in order to evaluate their performance in distinguishing those who need in-flight oxygen supplementation from those who do not.

## Materials and Methods

CF-patients, followed in our unit, aged over 18 years old, with stable disease, who intent to fly in the future were recruited. Thirty CF-patients were included in our study. To recruit healthy adults, an open call was launched by e-mail among medical students and young medical doctors. The participants had to fill a questionnaire with known diseases, current chronic medication, symptoms during previous flights and intention to fly in future. Individuals aged under 18 years old, with previous known diseases, symptoms during previous flights or with no intention to fly were excluded. We randomly selected 58 healthy adults. For both groups, we recorded age, sex, weight, height, body mass index (BMI) and smoking habits. Both spirometry and HAST was performed in CF-patients in a period of clinical stability with no change in their usual medication.

Spirometry breathing room air at sea level (Lisbon, 2 m above sea level, hereafter referred to as sea level) was performed before HAST using body screen from Viasys (Conshohocken, Pennsylvania, US) and the reference equation of Quanjer et al.^18^ to determine forced expiratory volume in 1 second (FEV_1_), Forced vital capacity (FVC) and calculate FEV_1_/FVC.

The normobaric HAST was performed at sea level and according to the recommendations of the BTS^4^ and Vohra and Klocke method^19^. Briefly, following this protocol, participants had to breath a FiO_2_ of 15% using a gas mixture with a supply of 99.993% nitrogen (Linde Healthcare, Lisboa, Portugal) through a 40% flow Venturi mask (Intersurgical, Berkshire, UK). Cardiorespiratory monitoring was performed using a polygraph and an oximeter in a hand finger (Alice PDX; Philips-Respironics, Murrysville, PA, USA; Nonin Medical, Plymouth, MN, USA). Parameters monitored included oxygen peripheral saturation, arterial pressure (Classic Check, Pic solution, Artsana, Grandate, Italy) and electrocardiography. As established by BTS, recommended HCT duration is between 20 and 25 min. Immediately before and 30 min after provocation an ABG sample was drawn and immediately analyzed (Rapid point 500, Siemens, Erlangen, Germany), to measure PaO_2ground_, oxygen saturation at ground (SatO_2ground_), PaO_2HALT_ and oxygen saturation after HAST (SatO_2HAST_). HAST results were then compared with the predictions made by the following equations:

- 1^st^ equation (Dillard et al.)^14^: PaO_2alt_= 0,410 x PaO_2ground_ + 17,652;
- 2^nd^ equation (Dillard et al.)^14^: PaO_2alt_= 0,519 x PaO_2ground_ + 11,855 x FEV_1_ (L) − 1,760;
- 3^rd^ equation (Dillard et al.)^14^: PaO_2alt_= 0,453 x PaO_2ground_ + 0,386 x FEV_1_ (%) + 2,44;
- 4^th^ equation (Gong et al.)^16^: PaO_2alt_= 0,88 + 0,68 x PaO_2ground_;
- 5^th^ equation (Kamin et al.)^17^: PaO_2alt_= PaO_2ground_ − 26,6.

## Statistical analysis

Clinical characteristic, spirometry and ABG results were analyzed and compared in the healthy controls and CF patients. Normal distribution of the data was assessed using Shapiro-Wilk test; Student’s *t-*test was performed for data with a normal distribution, and the non-parametric Wilcoxon test when that was not the case. The Chi-square test was preferred for categorical variables. To verify the effectiveness of the five equations in separating who needs oxygen during the flight from those who do not, receiving operator characteristic (ROC) curves were performed and the area under the curve (AUC) was calculated for each equation. This study was approved by the Research Ethics Committee of Centro Hospitalar Lisboa Norte (Lisboa, Portugal).

## Results

Out of the 88 randomly selected adults, all were caucasians and the majority were women (52.27%) with a median age of 22 years old. Between CF-patients and the healthy controls, there was a significant difference in weight, height and body mass index (BMI) (*p*<0.001), with CF-patients being thinner and shorter. In the control group, 18.96% were former smokers and 12.06% current smokers, while all of the CF-patients were non-smoker. No statistically significant difference was found for age and gender. Clinical characteristics of the sample studied are summarized in Table 1.

**Table 1.** Clinical characteristics of the sample studied.

| Characteristics | Total<br>(n=88) | CF patients<br>(n=30) | Healthy controls<br>(n=58) | p-value |
| --- | --- | --- | --- | --- |
| Age, years<sup>†</sup> | 22 [21; 24] | 23 [21.25; 30.75] | 22 [21; 23] | 0.099 |
| Gender - Men, n (%) | 42 (47.72) | 14 (46.66) | 28 (48.27) | 1 |
| Weight, kg<sup>†</sup> | 61 [53.75; 70.5] | 53 [50; 61.5] | 65 [57.25; 75] | <0.001 |
| Height, cm<sup>*</sup> | 167.52 ± 8.94 | 163.3 ± 7.02 | 169.71 ± 9.08 | <0.001 |
| BMI, kg/m<sup>2†</sup> | 22.23 [19.84; 23.87] | 19.93 [18.99; 21.92] | 23.08 [21.81; 23.25] | <0.001 |
| Non-smokers, n (%) | 70 (79.54) | 30 (100) | 40 (69.96) | <0.001 |

\*: mean ± standard deviation; †: median [interquartile range]; BMI: Body mass index.

We found significant differences when comparing the recorded functional respiratory values: CF-patients had lower FVC, FEV_1_ and FEV_1_/FVC (*p*< 0.001). CF-patients had also lower PaO_2ground_, SatO_2ground_, PaO_2HALT_, SatO_2HAST_ and lower minimal oxygen saturation (SpO_2_min) measured during HAST (*p*<0.001). Respiratory functional results are summarized in Table 2.

**Table 2.** Respiratory functional results of the sample studied.

| Results | Total<br>(n=88) | CF patients<br>(n=30) | Healthy controls<br>(n=58) | p-value |
| --- | --- | --- | --- | --- |
| FVC, L<sup>†</sup> | 3.74 [3.09; 4.75] | 2.76 [2.35; 3.23] | 4.21 [3.62; 5.23] | <0.001 |
| FVC, %<sup>†</sup> | 95.50 [81.5; 104] | 72.5 [60.5; 82] | 101 [94.25; 108] | <0.001 |
| FEV<sub>1</sub>, L<sup>†</sup> | 3.32 [2.23; 4.04] | 1.7 [1.18; 2.5] | 3.66 [3.28; 4.42] | <0.001 |
| FEV<sub>1</sub>, %<sup>†</sup> | 93 [71.75; 104] | 56 [37.25; 71.75] | 103 [93; 208.75] | <0.001 |
| FEV<sub>1</sub>/FVC, %<sup>†*</sup> | 84.9 [72.77; 88.26] | 63.3 [53; 73.13] | 86.58 [82.58; 91] | <0.001 |
| PaO<sub>2ground</sub>, mmHg<sup>†</sup> | 95.7 [85.9; 99.88] | 80.25 [71.68; 90.75] | 98.2 [94.98; 109.1] | <0.001 |
| PaO<sub>2HAST</sub>, mmHg<sup>†</sup> | 66.7 [59.13; 70.9] | 53.45 [45.9; 59.38] | 69.35 [65.48; 73.35] | <0.001 |
| SatO<sub>2ground</sub>, %<sup>†</sup> | 97.5 [96.7; 97.73] | 96.05 [94.35; 97] | 97.65 [97.4; 97.9] | <0.001 |
| SatO<sub>2HAST</sub>, %<sup>†</sup> | 93.5 [91.35; 97.8] | 89.15 [83.85; 91.7] | 94.4 [93.13; 95.15] | <0.001 |
| SpO<sub>2min</sub>, %<sup>†</sup> | 88 [85; 93] | 83 [80; 85.75] | 88.5 [87.25; 90.75] | <0.001 |

†: median [interquartile range]; \*: using the reference equation of Quanjer et al.<sup>18</sup>; FVC: Forced volume capacity; L: Liters; FEV<sub>1</sub>: Forced expiratory volume in 1 second; PaO<sub>2ground</sub>: partial arterial oxygen pressure measured at ground; PaO<sub>2HAST</sub>: partial arterial oxygen pressure measured at the end of the hypoxia challenge test; SatO<sub>2ground</sub>: oxygen saturation at ground measured by arterial blood gas; SatO<sub>2HAST</sub>: oxygen saturation at the end of the hypoxia challenge test measured by arterial blood gas; SpO<sub>2min</sub>: lower minimal oxygen saturation measured during the hypoxia challenge test measured by finger probe.

None of the healthy controls reached a PaO_2HAST_ below 50 mmHg, while 11 (36.7%) of the CF-patients had lower values, with 42.7 mmHg being the lowest founded value – Table 3.

**Table 3.** Mean PaO_2_ at altitude (mmHg) by HAST and predicted the five equations.

|  | **Total**<br>**(n=88)** | **CF patients**<br>**(n=30)** | **Healthy controls**<br>**(n=58)** | **p-value** |
| --- | --- | --- | --- | --- |
| **HAST** | 64.78 ± 11.18 | 54.31 ± 9.32 | 70.59 ± 7.21 | **<0.001** |
| Lowest value | 42.70 | 42.70 | 59.20 |  |
| **1<sup>st</sup> equation\*** | 55.45 ± 4.74 | 50.81 ± 4.69 | 57.85 ± 2.44 | **<0.001** |
| Lowest value | 42.58 | 42.58 | 51.11 |  |
| **2<sup>nd</sup> equation\*** | 83.74 ± 19.21 | 63.06 ± 16.34 | 94.44 ± 9.28 | **<0.001** |
| Lowest value | 37.58 | 37.58 | 76.54 |  |
| **3<sup>rd</sup> equation\*** | 77.46 ± 15.22 | 60.38 ± 13.54 | 86.29 ± 5.24 | **<0.001** |
| Lowest value | 38.75 | 38.75 | 74.14 |  |
| **4<sup>th</sup> equation\*** | 63.57 ± 7.87 | 55.87 ± 7.77 | 67.55 ± 4.05 | **<0.001** |
| Lowest value | 42.22 | 42.22 | 56.37 |  |
| **5<sup>th</sup> equation\*** | 65.59 ± 11.57 | 54.27 ± 11.43 | 71.44 ± 5.96 | **<0.001** |
| Lowest value | 34.20 | 34.20 | 55 |  |

When CF-patients were divided by PaO_2HAST_ (below or above 50 mmHg), a significant difference was found for weight (*p*=0.002), BMI (*p*<0.002), FVC, FEV_1_ (*p*<0.001), FEV_1_/FVC (*p*=0.003), PaO_2ground_ (*p*=0.008) and SatO_2ground_ (*p*<0.007), with patients with PaO_2HAST_ <50 mmHg having lower values. No difference was found for age (*p*=1), gender (*p*=0.466) or height (*p*=0.219) between CF-patients.

## Adjustment between groups

As the healthy controls and CF-patients have significant differences, we performed an adjustment for tobacco usage, BMI and height. All smokers and former smokers were excluded. Out of the 70 non-smoker adults (40 healthy controls and 30 CF-patients), the majority were women (54.28%) with a median age of 22 years old. Between CF-patients and the healthy controls, as expected, there was a significant difference in weight (*p*< 0.001), height (*p*=0.003) and BMI (*p*< 0.001). No statistically significant difference was found for age (*p*= 0.092) and gender (p= 1).

Significant differences were found when analyzing functional respiratory values: CF-patients had lower FVC, FEV_1_ and FEV_1_/FVC (*p*< 0.001). CF-patients had also lower PaO_2ground_, SatO_2ground_, PaO_2HAST_, SatO_2HAST_ (*p*< 0.001) and a lower SpO_2_min measured during HAST (*p*< 0.001).

With all five equations, the mean predicted PaO_2alt_ had a significant difference between healthy controls and CF-patients (*p*<0.001). When the five equations were compared to each other, the mean predicted PaO_2alt_ had also a significant difference (*p*≤0.005), but when a cut-off of 50 mmHg was used, no difference was found (*p*=0,51).

To prevent a confounding effect by BMI and height, we used a correction based on multiple linear regression. This method allows to take into consideration the variability associated with each factor. A significant difference was still found between the groups on FVC, FEV_1_ and FEV_1_/FVC, PaO_2ground_, SatO_2ground_, PaO_2HAST_, SatO_2HAST_, SpO_2_min and predicted PaO_2alt_ by the five equations (*p*< 0.001).

## Equation comparison

### Healthy adults and CF-patients

When the values of PaO_2alt_ in the healthy controls and CF-patients were compared, a significant difference was found in all five equations (*p*< 0.001). The lowest predicted PaO_2alt_ was found with the 5^th^ equation, on the CF-patient group (34.2 mmHg) – Table 3. This value was lower than the respective PaO_2HAST_ found. When a cut-off of 50 mmHg for PaO_2alt_ was used, none of the control had a predicted PaO_2alt_ < 50 mmHg. We used a non-parametrical approach for all variables when CF-patients were divided by PaO_2HAST_ (below or above 50 mmHg), because the groups became smaller. In this case, a significant difference was found for weight (*p*=0.002), BMI (*p*<0.002), FVC, FEV_1_ (*p*<0.001), FEV_1_/FVC (*p*=0.003), PaO_2ground_ (*p*=0.008) and SatO_2ground_ (*p*<0.007). Patients with PaO_2HAST_ <50 mmHg had the lowest values. No difference was found on age (*p*=1), gender (*p*=0.466) or height (*p*=0.219) between CF-patients.

Using this cut-off, there were no evidences of significant differences between the case distribution among the equations (*p=*0.369). However, when we compare with the case distribution based on the results from the HAST test, some CF-patients were misclassified – Table 4 and Table 5.

**Table 4.**
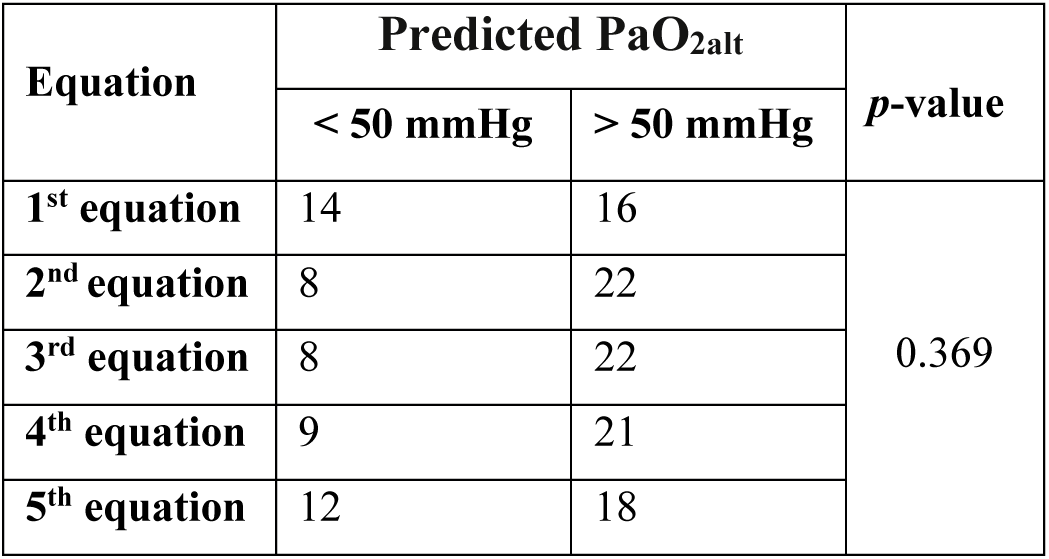
Division of CF-patients by predicted PaO_2_ at altitude (PaO_2alt_, mmHg) by the five equations.

| **Equation** | **Predicted PaO<sub>2alt</sub>** |  | **p-value** |
| --- | --- | --- | --- |
|  | **< 50 mmHg** | **> 50 mmHg** |  |
| **1<sup>st</sup> equation** | 14 | 16 | **0.369** |
| **2<sup>nd</sup> equation** | 8 | 22 |  |
| **3<sup>rd</sup> equation** | 8 | 22 |  |
| **4<sup>th</sup> equation** | 9 | 21 |  |
| **5<sup>th</sup> equation** | 12 | 18 |  |

**Table 5.** Division of CF-patients by HAST results and by predicted PaO_2_ at altitude (PaO_2alt_, mmHg) by the five equations.

|  |  | **1<sup>st</sup> equation** |  | **2<sup>nd</sup> equation** |  | **3<sup>rd</sup> equation** |  | **4<sup>th</sup> equation** |  | **5<sup>th</sup> equation** |  |
| --- | --- | --- | --- | --- | --- | --- | --- | --- | --- | --- | --- |
|  |  | **<50**<br>**mmHg** | **>50**<br>**mmHg** | **<50**<br>**mmHg** | **>50**<br>**mmHg** | **<50**<br>**mmHg** | **>50**<br>**mmHg** | **<50**<br>**mmHg** | **>50**<br>**mmHg** | **<50**<br>**mmHg** | **>50**<br>**mmHg** |
| **HAST**<br>**results** | **< 50**<br>**mmHg** | 7 | 4 | 7 | 4 | 7 | 4 | 7 | 4 | 7 | 4 |
|  | **> 50**<br>**mmHg** | 7 | 12 | 1 | 18 | 1 | 18 | 2 | 17 | 5 | 14 |

We found no statistically significant difference on the sensibility of each equation (63.64%) and the specificity was 63.16% (1^st^ equation), 94.74% (2^nd^ and 3^rd^), 89.47% (4^th^) and 74.68% (5^th^). The positive predictive value was 50% (1^st^ equation), 87.5% (2^nd^ and 3^rd^), 77.78% (4^th^) and 58.33% (5^th^). The negative predictive value was 75% (1^st^ equation), 81.82% (2^nd^ and 3^rd^), 80.95% (4^th^) and 77.78% (5^th^).

In order to compare the performance of the five equations with the cut-off of 50 mmHg, ROC curves were performed (Figure 1). The 3^rd^ equation had the best performance with an AUC of 88.04%.

**Figure 1.**
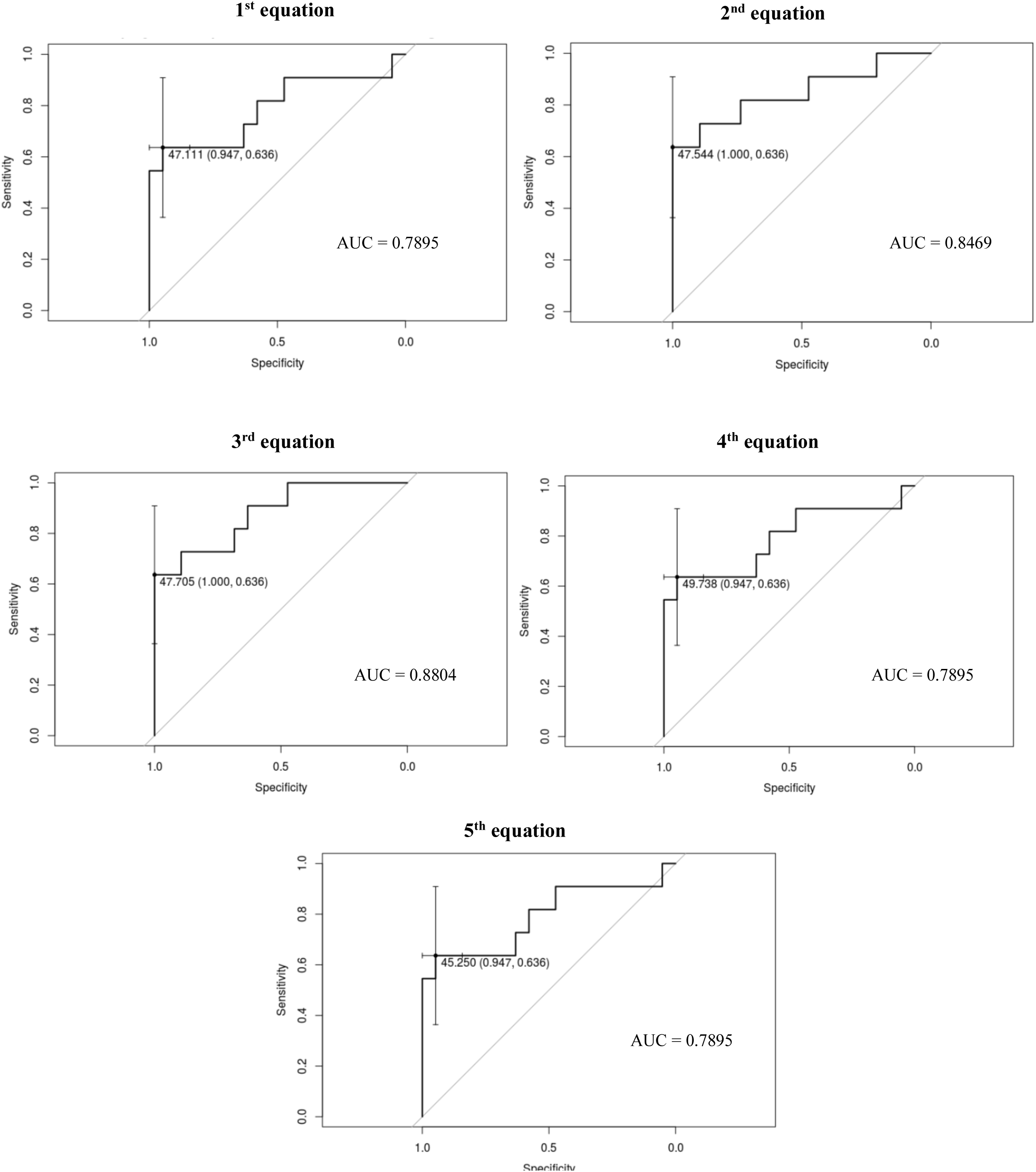
Receiving Operator Characteristic (ROC) curves and Area Under the Curve (AUC) of the five equations analyzed.

## Prediction of in-flight oxygen need in CF-patients

To test whether the requirement for in-flight oxygen supplementation could be predicted by PaO_2ground_ and/or by FEV_1_ (in Liters and predicted percentage), we analyzed the CF-patients with PaO_2HAST_ < 50 mmHg (n=11). Of this sub-group eight (72.73%) had FEV_1_ < 50%, six (54.55%) had PaO_2ground_ < 70 mmHg, two of which had < 65 mmHg.

A relation between FEV_1(in liters)_/ PaO_2HAST_, FEV_1(%)_/ PaO_2HAST_ and PaO_2ground_/ PaO_2HAST_ was found (*p*< 0.001) – Figure 2.

**Figure 2.**
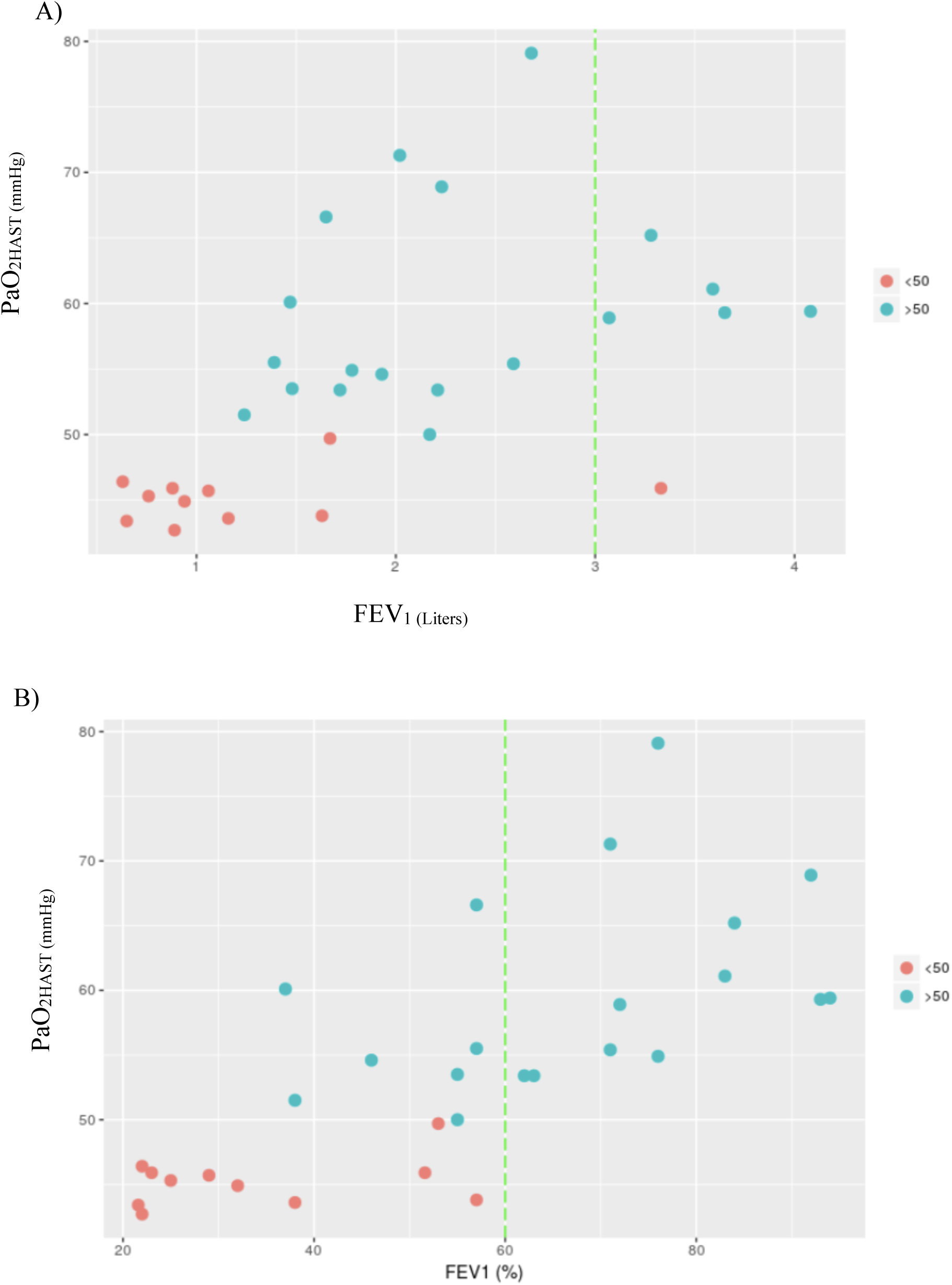
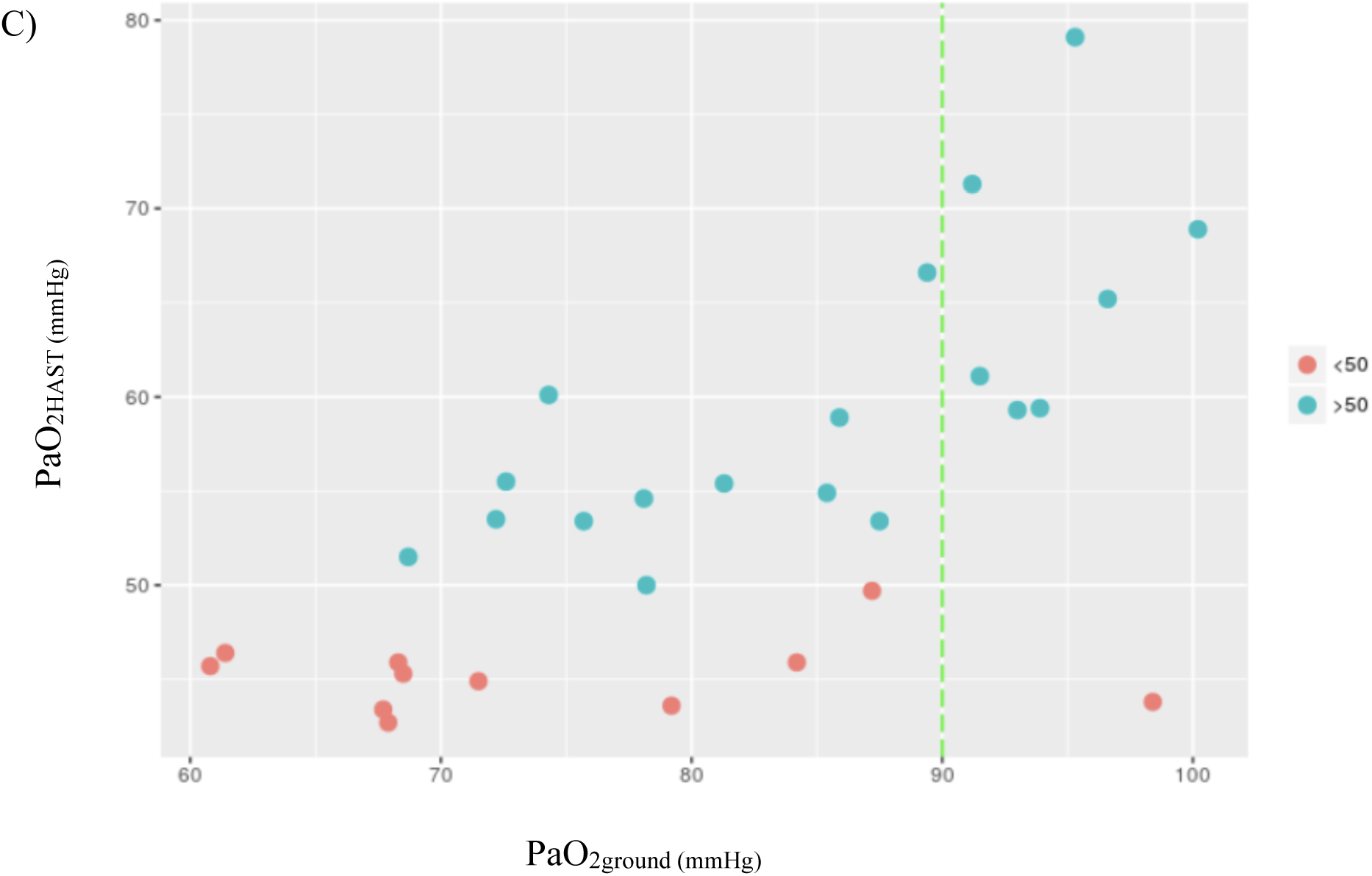
Relation between: (A) FEV_1(in liters)_/ PaO_2HAST_; (B) FEV_1(%)_/ PaO_2HAST_; (C) PaO_2ground_/ PaO_2HAST_.

Out of the patients that required in-flight oxygen supplementation, 10 had FEV_1_ < 2 Liters, one FEV_1_ > 3 Liters, all had FEV_1_ < 60 % of the predicted value and PaO_2ground_ > 60 mmHg, with one patient having PaO_2ground_ > 90 mmHg.

## Discussion

Guidelines refer to predictive equations of altitude hypoxia as either a screening tool for recommending a HAST or as a replacement when this test is not available,^11^ but the accuracy and predictive value of the available equation in relation to the HAST *Gold Standard* has not been previously evaluated in CF patients in comparison to healthy individuals. In our sample, we found a significant difference between healthy and CF-patients even when adjusted for tobacco usage, BMI or height. In our sample, 11 CF-patients and none of the healthy controls reached a PaO_2HAST_ below 50 mmHg. Patients who needed in-flight oxygen supplementation were thinner, had lower FVC, FEV_1_, FEV_1_/FVC, PaO_2ground_ and SatO_2ground_ than those who did not.

While the 3^rd^ equation performed better than the other four, when compared to HAST results, none was able to correctly identify CF-patients that required in-flight oxygen supplementation. In a previous study, Martin et al.5 tested the prediction of the first four equations in 15 adults with CF and concluded that the four equations overestimated the need of in-flight oxygen. In our sample, only the first and fifth equations overestimated the HAST results. While overestimation will lead to unnecessary prescription of oxygen in flight with no clinical risk but increasing flight cost, possibly making the trip unfeasible, the opposite could lead a significant clinical risk. Therefore, we do not recommend to use these equations as screening tools or substitute of HAST.

According to some literature,^3,12^ CF-patients with FEV_1_ > 50% or PaO_2ground_ > 60 mmHg can safely travel without oxygen supplementation. Therefore, this recommendation would exempt the performance of HAST in these patients. In fact, in our sample, FEV_1_ and PaO_2ground_ correlated with HAST results, with FEV_1_ in predicted percentage being the most accurate. However, contrary to the recommendations described above, all patients that needed in-flight oxygen had a FEV_1_ < 60% while no other cut-off on FEV_1_ in Liters or PaO_2ground_ was able to correctly predict those who reached PaO_2HAST_ below 50 mmHg. We do not recommend to use the previous values of FEV_1_ (> 50% of the predicted value) or PaO_2ground_ (> 60 mmHg) as, in our sample, tree patients with FEV_1_ > 50% and all 11 patients that needed in-flight oxygen had PaO_2ground_ > 60 mmHg.

We found a large variability between the results given by HAST and the values predicted by the different equations studied. The third equation produced predictions closer to the results observed by HAST, but given the individual difference found, the authors recommend to, whenever possible, perform a HAST to analyze commercial air flights risks.

## Acknowledgements

The authors are grateful to the staff of Serviço de Pneumologia, Departamento do Tórax, 1649-035 Lisboa, Portugal for technical support. No specific grant from funding agencies in the public, commercial, or not-for-profit sectors was used to support this research.

